# ASIC1a is required for neuronal activation via low-intensity ultrasound stimulation in mouse brain

**DOI:** 10.1101/2020.07.10.196634

**Authors:** Jormay Lim, Ya-Cherng Chu, Chen-Ming Hao, Wei-Hao Liao, Shao-Shien Lin, Sherry Hsu, Hsiao-Hsin Tai, Ya-Chih Chien, Dar-Ming Lai, Wen-Shiang Chen, Chih-Cheng Chen, Jaw-Lin Wang

## Abstract

Accumulating evidence has shown transcranial low-intensity ultrasound can be potentially a non-invasive neural modulation tool to treat brain diseases. However, the underlying mechanism remains elusive, because the majority of studies on animal models applying rather high-intensity ultrasound that cannot be safely used in humans. Here we showed low-intensity ultrasound was able to activate neurons in the mouse brain and repeated ultrasound stimulation resulted in adult neurogenesis in specific brain regions. *In vitro* calcium imaging studies showed that a specific ultrasound stimulation mode, which combined with both ultrasound-induced pressure and acoustic streaming mechanotransduction, is required to activate cultured cortical neurons. ASIC1a and the tether-mode mechanotransduction were involved in the low-intensity ultrasound-mediated mechanotransduction and cultured neuron activation, which was inhibited by ASIC1a blockade and cytoskeleton-modified agents. In contrast, the inhibition of mechanical sensitive channels involved in bilayer-model mechanotransduction like Piezo or TRP proteins did not affect the ultrasound-mediated neuronal activation.

**SIGNIFICANCE:** CNS neurons have no sensory function, protected by the skull. For this reason, brain neuromodulation by ultrasound were either done at a high intensity or through auditory nerves. We demonstrate in this study CNS neurons react to ultrasound stimulation at an intensity (5 mW/cm^2^) far lower than typical therapeutic ultrasound (>30 mW/cm^2^). Using micropipette ultrasound in calcium imaging, we show the reactions of CNS neurons to ultrasound is through ASIC1a channels, pointing to the molecular basis for direct ultrasound neuromodulation at low intensity. Furthermore, we also show evidence of neurogenesis with the same ultrasound stimulation, suggesting potential therapeutic translation.

## Introduction

Transcranial ultrasound such as opening blood-brain barrier (BBB) (1) for localized drug release and modulating neural activity (2-4) has been used for therapeutic treatments of various brain diseases. Many *in vivo* animal experiments and human clinical trials (*SI Appendix*, Table 1) proved the clinical potential of transcranial ultrasound stimulation. With the increased interest of this technique, the mechanisms underlying ultrasound-mediated neural modulation has also recently been learned. A study showed high-intensity transcranial ultrasound can elicit a startle-like motor response via an indirect auditory mechanism (5). Emerging sonogenetics in worm model also identified and engineered TRP-4 channels as a sensor for the ultrasound stimulus to activate neurons in living organisms at pressure level above 0.5 MPa (6), and ultrasound at 0.1 MPa acoustic pressure was found to activate neurons via Piezo 1 mechanosensitive ion channel (7). Nonetheless, the energy intensity or acoustic pressure of most clinical trials or basic researches used for BBB opening or neuromodulation are both high, and safety issue of this technique in clinical application remains a concern.

In this study, a much lower intensity ultrasound at the order lower than 10 mW/cm^2^ is proposed to activate neurons via mechanosensitive ion channels in mammals’ brain for potential clinical application. Mechanosensitve ion channels are mainly grouped into bilayer model, such as PIEZO and TRP channels gated by membrane tension change, and extracellular matrix tethered model such as acid sensing ion channels (ASICs) (8-10). Here we aim to identify possible mechanical sensors in mouse brain that can directly respond to low-intensity ultrasound.

## Results

### Transcranial ultrasound induced p-ERK in the cortex, hippocampus and amygdala of mouse brain

We kept ultrasound exposure to below 10 mW/cm^2^ (I_SATA_) in our experiments to ensure safe therapeutic applicability. The phosphorylation of extracellular-signal-regulated kinase (p-ERK), an established indicator of immediate neuronal activation (11), was used to evaluate whether transcranial low-intensity ultrasound can stimulate neuronal activity in mouse brain. Mice with 1-minute ultrasound exposure (Fig. 1 *A*) had shown significant increase of p-ERK positive cells in certain brain regions, such as the cortex (Fig. 1 *B-D*), hippocampus (Fig. 1 *E-G*), and amygdala (Fig. 1 *H-J*) as compared with those received sham treatment (*SI Appendix*, Table 2). More specifically, increased p-ERK expression generally occurred upon ultrasound stimulation in the visual, somatosensory, auditory, temporal associations, retrosplenial, piriform, and entorhinal areas of mouse cortex (*SI Appendix*, Fig. S1 and S2). In hippocampus, CA1 and CA2 were dramatically lightened up with p-ERK signals in response to ultrasound whereas CA3 and dentate gyrus showed sparsely stimulated (*SI Appendix*, Fig. S1 and S2). In amygdala, the central amygdala nucleus showed the strongest p-ERK signals, while medial and basolateral also obviously increased in p-ERK signals (*SI Appendix*, Fig. S1 and S2). We also observed a consistently unchanged p-ERK staining in the paraventricular nucleus of hypothalamus (PVH) (*SI Appendix*, Fig. S3), revealing an intriguing regional specificity of the ultrasound response.

**Fig. 1.**
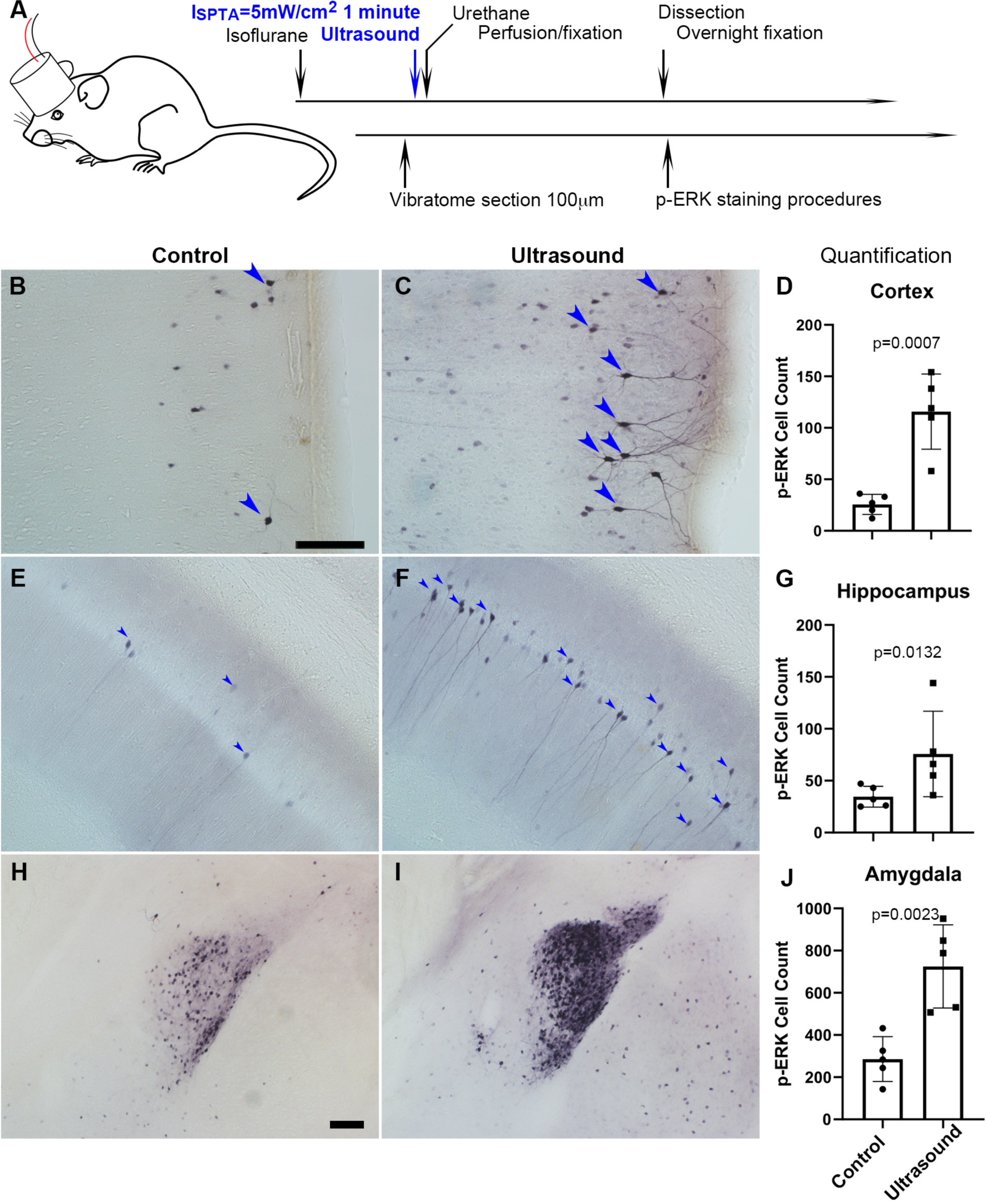
Transcranial ultrasound induces pERK expression in neurons of the cortex, hippocampus and amygdala of mouse brain. (*A*) Illustration depicting mouse head stimulated by 1 MHz transducer which was positioned in between the mouse nasal process of maxilla and the axis of mouse ear of an anesthetized mouse. (*B*) Micrograph representing cortical region with basal level of p-ERK staining in sham control mice (n=5), scale bar 100 μm. Sham control mice were handled with similar procedures of placing transducer on the head without turning on the ultrasound function generator. (*C*) Micrograph representing cortical region with p-ERK staining stimulated by ultrasound (I_SPTA_=5mW/cm_2_, 1 minute) (n=5). (*D*) Quantitative bar graph of the number of p-ERK stained cells within comparable area of 1.224mm^2^ (Length 1275μm, Width 960μm), showing significant difference (p=0.0007) of cell count by ImageJ. (*E*) Micrograph representing hippocampal region with basal level of p-ERK staining in sham control mice. (*F*) Micrograph representing hippocampal region with p-ERK staining in mice stimulated by ultrasound. (*G*) Bar graph showing quantification of significantly p-ERK different cell count (1.224mm^2^) (p=0.0132) in hippocampal region. (*H*) Micrograph representing amygdala of sham controls. Scale bar 100 μm (*I*) Micrograph representing amygdala of ultrasound stimulated mice. (*J*) Quantification of amygdaloid significant difference (p=0.0023) of p-ERK cell count (1.224mm^2^).

### Micropipette guided ultrasound as the mechanical stimuli with combined ultrasound and acoustic streaming

To determine whether low-intensity ultrasound can directly activate neurons mechanically, we next used in vitro calcium-imaging approach to probe the possible ion channels responding to ultrasound mechanical stimulus in cultured cortical neurons. A micropipette was used to guide ultrasound to cultured cells (Fig. 2 *A*). The device can generate either an ultrasound induced pressure predominant condition (2000 mVpp, DF 0.05%, measured peak pressure versus the distance to the pipette tip shown in *SI Appendix*, Fig. S4 *A*) or an acoustic streaming predominant condition (100 mVpp, DF 100%, flow pattern for the acoustic streaming shown in *SI Appendix*, Fig. S4 *A* and Movie S1), or a mixed loading condition (700 mVpp, DF 20%) depending on duty cycle applied. Ultrasound pressure predominant conditions (up to 2000 mVpp, DF 0.05%) generated compressional stress on cells that were not able to elevate calcium responses (Fig. 2 *B* and *SI Appendix*, Movie S2), while acoustic streaming predominant conditions (up to 100 mVpp, DF 100%) invoked shear stress that could only activate little calcium responses in neurons in the maximal energy level applied (Fig. 2 *B*); whereas a mixed loading condition (700 mVpp, 20% DF) effectively yielded much higher calcium responses (Fig. 2 *B* and *SI Appendix*, Movie S3). The ultrasound predominant mode with only compressional stress cannot induce a response even when the stimulation was extended to 10 seconds (*SI Appendix*, Fig. S5 *A*). Similarly, the prolong stimulation of acoustic streaming invoking shear stresses also only elevated mildly the amplitude of calcium response (*SI Appendix*, Fig. S5 *B*). On the other hand, the combined compression stress with acoustic streaming can reproducibly elevated calcium response even in 1.5 second stimulation (Fig. 2 *B* and *SI Appendix*, Movie S3, Fig. S5 *C*). The ultrasound-induced calcium responses were dose-dependent with a threshold of 400 mVpp (8 kPa) and EC_50_ of 700 mVpp (12kPa) (Fig. 2 *C* and *SI Appendix* Fig. S5 *D-K*). The corresponding stress levels of ultrasound at 400 mVpp, 500 mVpp, 700 mVpp or 900 mVpp were 8 kPa, 8.72 kPa, 12 kPa and 15.3 kPa, respectively.

**Fig. 2.**
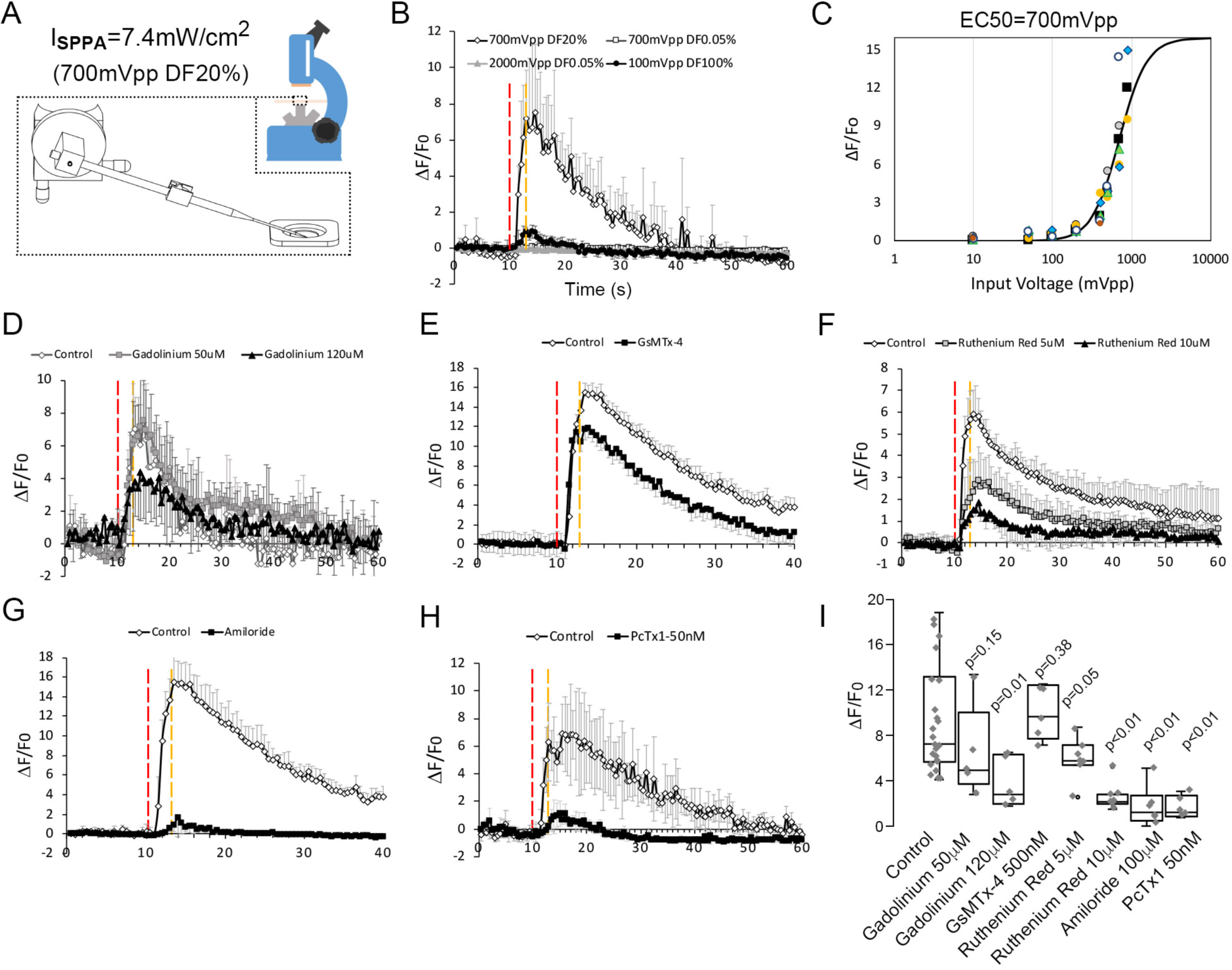
Neuronal calcium signals induced by micropipette-guided ultrasound suppressed by ASIC1a inhibitors. (*A*) A micropipette positioned to the cortical neurons cultured on a 30mm cover slips mounted to a chamber coupled to microscope platform. Calcium signals recorded from neurons stained by Invitrogen™ Oregon Green™ 488 BAPTA-1, AM cell permeant. (*B*) Line graphs of averaged calcium signals in 4 neurons stimulated by micropipette ultrasound for 3 seconds with an input voltage 2000mVpp, duty factor (DF) 0.05%; or 100mVpp, DF100% or 700mVpp, DF20%. The red-dotted line denotes start of the stimulation while the yellow-dotted line denotes the end. (*C*) Calcium responses as a function of micropipette ultrasound in 20%DF. Dose-dependent (input voltages from 10mVpp, to 900mVpp, DF20%) responses of ultrasound with an EC50 of 700mVpp is shown (n=5). (*D*) Effects of gadolinium (120 μM), a non-selective blocker of mechanically sensitive ion channels, on calcium signals in cortical neurons. (*E*) Effects of GsMTx-4 (500nM), a selective Piezo inhibitor, on calcium signals in cortical neurons. (*F*) Effects of Ruthenium red (10 μM), a non-selective TRP inhibitor, on calcium signals in cortical neurons. (*G*) Effects of amiloride (100 μm), an ASICs family inhibitor, on calcium signals in cortical neurons. (*H*) Effects of PcTx1 treated (50 nM), a selective ASIC1a inhibitor, on calcium signals in cortical neurons. (*I*) Statistical analyses of channel blockers on ultrasound-induced calcium signals in cortical neurons.

### Neuronal calcium signal upon ultrasound simulation suppressed by ASIC channels inhibitors

The micropipette ultrasound mechanotransduction was pharmacologically tested with selective or non-selective blockers of mechanosensitive ion channels for Piezo, TRP, and ASICs. First, the treatment with Gadolinium (10 μM), a non-selective blocker of mechanically sensitive ion channels, partially suppressed the ultrasound-induced calcium signals (Fig. 2 *D*). Treating the cells with a selective Piezo inhibitor, GsMTx-4, led to a marginal (not significant) inhibition of the ultrasound-induced calcium elevation as compared with the vehicle control (Fig. 2 *E*). Instead, the treatment with a TRP blocker ruthenium red (1-10 μM) partially suppressed the calcium signals (Fig. 2 *F*), whereas the non-selective ASIC inhibitor, amiloride, totally abolished the calcium signals by micropipette ultrasound (Fig. 2 *G*). Above results suggested ASICs might be the major channels involved in micropipette ultrasound mechanotransduction. To narrow down the specific candidate of ASICs, ASIC1a inhibitor, PcTx1 (50nM) was tested. PcTx1 (50 nM) significantly inhibited the calcium response by micropipette ultrasound (Fig. 2 *H* and *SI Appendix*, Fig. S5 *C*). The relative inhibitions of above channel blockers were summarized in Fig. 2*I*. We further tested the dose-dependent inhibition of PcTx1 on micropipette ultrasound and determined an IC50 of 0.2nM, suggesting a homotrimeric ASIC1a is the mechanoreceptor in action (Fig. 3 *A* and *SI Appendix*, Fig. S6 *A-F*). To validate how ASIC1a activation would lead to calcium responses to ultrasound, we treated the cells with calcium chelating agent, ethylene glycol-bis(β-aminoethyl ether)-N,N,N’,N’-tetraacetic acid (EGTA) (1-5mM) to block the extracellular calcium. The results showed that calcium influx was absolutely essential (Fig. 3 *B* and *SI Appendix*, Fig. S6 *G* and *H*). To examine whether endoplasmic reticulum calcium was involved in calcium signaling, we found the calcium surge of cells treated with the RyR inhibitor, JTV519 fumarate (10 μM) (*SI Appendix*, Fig. S6 *G* and *I*) was partially inhibited, while as the IP3R inhibitor, (-)-Xestonspongin C (1 μM) was most inhibited (Fig. 3 *B*).

**Fig. 3.**
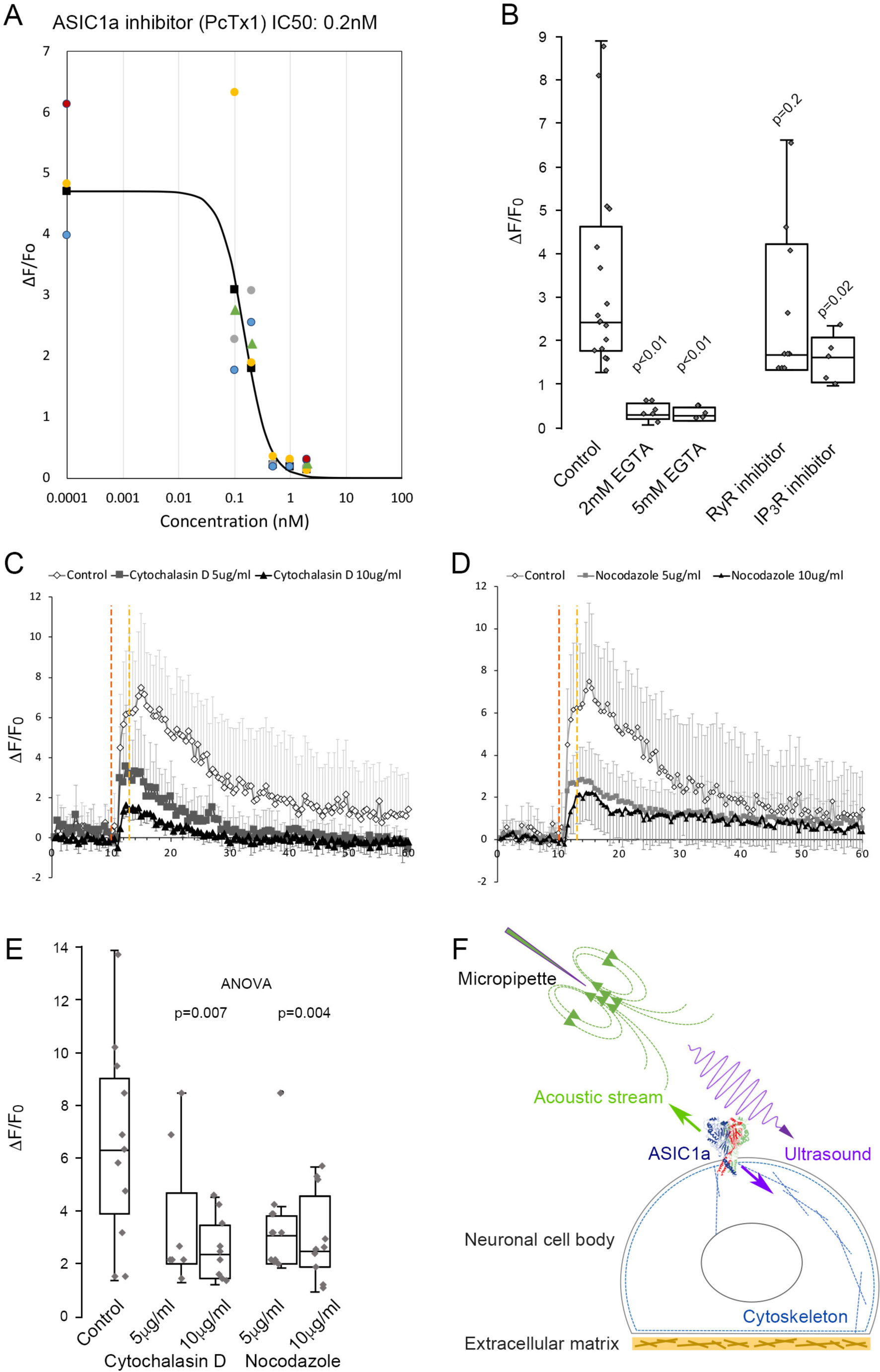
ASIC1a as a mechanoreceptor responsive to mechanical stimuli with combined ultrasound and acoustic streaming. (*A*) PcTx1 dose-dependent inhibition curve of calcium responses induced by micropipette ultrasound of 700mVpp, DF20% for 3-second. (*B*) Whisker plots showing comparison of peak ΔF/F_0_ within 3-5 seconds upon micropipette ultrasound (700mVpp, DF20%, 3-second) stimulation in the untreated control primary neurons, 2- or 5mM EGTA treated neurons, RyR inhibitor JTV519 fumarate (10 μM) treated neurons, or IP_3_R inhibitor (-)-Xestonspongin C (1 μM) treated neurons. Student t-test with *p* value compared to control listed above the whisker plot. (*C*) Graph showing calcium response of actin polymerization inhibitor Cytochalasin D (5-10 μg/ml) treated neurons compared to untreated control. (*D*) Calcium signals showing the effect of the microtubule assembly inhibitor nocodazole (5-10 μg/ml) on neurons. (*E*) Whisker plots showing comparison of peak ΔF/F_0_ within 3-5 seconds upon micropipette ultrasound (700mVpp, DF20%, 3-second) stimulation in the untreated control primary neurons, cytochalasin D 5- or 10 μg/ml treated neurons, nocodazole 5- or 10 μg/ml treated neurons. Statistical *p* values of one-way ANOVA analysis were listed to show the significance of treatment. (*F*) Cartoon depicting ultrasound stimulating ASIC1a in the cell body of a neuron under the micropipette ultrasound stimulation. Green arrow represents the pulling force of acoustic stream and purple arrow represents the compression force of ultrasound that results in cytoskeletal rearrangement.

### ASIC1a mechano-response required cytoskeletal dynamics

Previous studies have shown ASICs are involved in tether mode mechanotransduction, which relies on intact cytoskeletal structures (8, 12). We likewise treated the cells with either actin polymerization inhibitor, cytochalasin D (5-10 μg/ml) or microtubule assembly inhibitor, nocodazole (5-10 μg/ml). Indeed, inhibition of cytoskeletal dynamics could dose dependently and significantly suppress the calcium response stimulated by micropipette ultrasound (Fig. 3 *C-E*). The collated data revealed a novel mode of ultrasound mechanotransduction with a combination of compression force and shear force that activates ASIC1a channels in mouse neurons via tether mode mechanotransduction (Fig. 3 *F*).

### Transcranial ultrasound treatments promoted neurogenesis in dentate gyrus

We next investigated whether the low-intensity ultrasound stimulation in mouse brain could lead to a favorable outcome in terms of adult neurogenesis. We selected doublecortin (DCX) as a marker for neurogenesis in dentate gyrus (13-15). Compared to the non-treated controls, after 3 consecutive days of 5 minute ultrasound treatments (5 mW/cm^2^), DCX staining in dentate gyrus at day 4 and day 7 showed a significant 2-fold increase (Fig. 4). The results indicated that repeated stimulation of low-intensity ultrasound on mouse brain might achieve beneficial neural modulation and lead to neurogenesis in dentate gyrus.

**Fig. 4.**
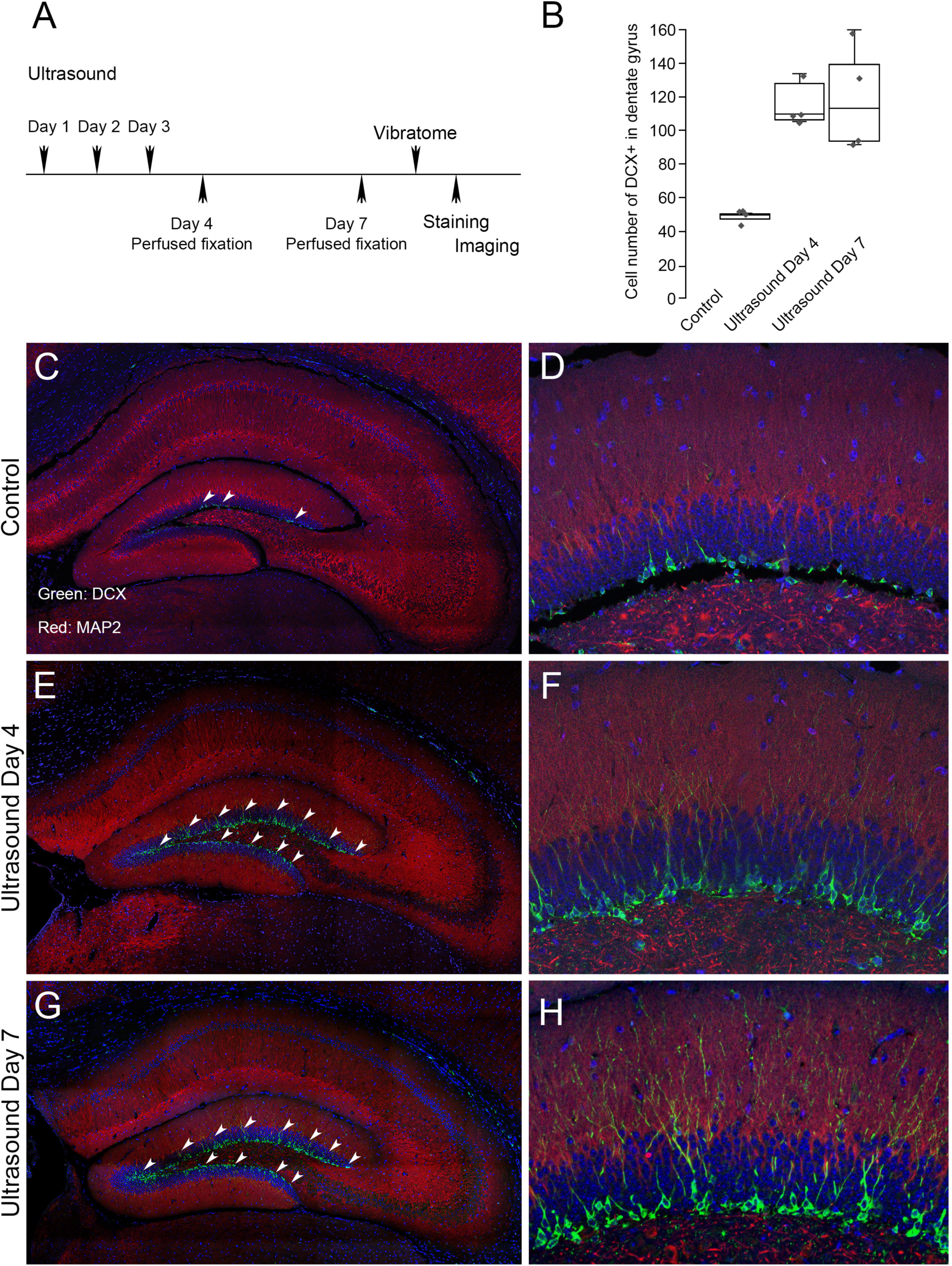
Neurogenesis in dentate gyrus induced by repeated transcranial ultrasound treatments. (*A*) The mice were treated 3 consecutive days by ultrasound of 4mW/cm^2^, 1% for 5 minutes, subsequently perfused fixed at day 4 or day 7 and brains were dissected from the head and sectioned for immunofluorescence procedures. The DCX staining in dentate gyrus of treated mice were compared to control untreated one. (*B*) Cell count with clear DAPI stained nucleus surrounded by DCX markers compared in control, day 4 and day 7 post-ultrasound treatments. Statistical analysis: p=0.0013 and F ratio=15.18 in one-way ANOVA (n=4). (*C, D*) Representative micrograph of untreated mice. The vibratome coronal brain sections (100 μm) of dentate gyrus region immunofluorescently stained for DCX (green) and MAP2 (red). Blue color indicates DAPI stained nuclei. Representative micrograph showing. (*E, F*) Representative micrograph from ultrasound treated mice fixed at day 4. (*G, H*) Representative micrograph from ultrasound treated mice fixed at day

## Discussion

Accumulating evidence has shown ASICs are involved in different types of mechanotransduction in the sensory nervous system, including nociception, barorecpetion, proprioception, and hearing (8, 16, 17). However, the mechanosensitive role of ASICs in the brain is still not known, although ASIC1a is a predominant acid sensor modulating neural activity in physiological and pathological conditions (18, 19). Here we demonstrated low-intensity ultrasound could modulate neural activity in mouse brain and directly activate neurons (Fig. 1) via an ASIC1a-depenent manner (Fig. 2-3). Moreover, repeated transcranial low-intensity ultrasound stimulations are safe and able to elicit adult neurogenesis in mouse brains (Fig. 4).

Although ASIC1a was determined as the molecular determinant involved in low-intensity ultrasound mechanotransduction, non-selective mechanosensor inhibitors such as gadolinium and ruthenium red were partially suppressing the calcium response triggered by micropipette ultrasound (Fig. 2 *G* and *H*). Of note gadolinium also blocked ASICs in μM to mM ranges. However, we cannot rule out a role of TRP channels in the micropipette ultrasound mechanotransduction, because there is no evidence showing ruthenium can also inhibit ASICs. Previous studies have proposed a role of TRP for ultrasound-mediated mechanotransduction while high-intensity ultrasound was applied. More studies are required to validate the role of TRP in low-intensity ultrasound mechanotransdcution or the unexpected role of ruthenium red in neuronal ASIC1a signaling pathways.

ASIC1a is widely expressed in the brain and could form as homotrimeric and heterotrimeric channels with different sensitivity to PcTx1 inhibition (20, 21). Specifically, heterotrimeric ASIC2b/ASIC1a can be inhibited 50% by approximately 3 nM (21) PcTx1 and ASIC1a/ASIC2a heterotrimeric can be inhibited by 50 nM(20) whereas 0.5 nM to 1 nM can inhibit the homotrimeric ASIC1a (22, 23). Therefore, since PcTx1 in low-doses effectively inhibited ASIC1a-mediated calcium signal by micropipette ultrasound, homotrimeric ASIC1a channels may be the predominant subtype involved in ultrasound mechanotransduction in cortical neurons (Fig. 3 *A* and *SI Appendix*, Fig. S6).

To explain how ultrasound could activate ASIC1a via the tether-mode mechanotransduction, we hypothesized a physical effect of micropipette ultrasound at cell level; which the acoustic streaming imposes shear stress on cell apical surfaces while ultrasound exerts compressional stresses throughout the cells. Considering the tether model *in vitro*, we argue, in a mixed loading condition (Fig. 2 *B* and 3 *F*), the extracellular domains of ASIC1a are under shear force pulling the protein to the flow direction while the intracellular domains of the ASIC1a are connected to cortical actin or other cytoskeleton, which experiences dynamic reorganization coupling with membrane withdrawals in response to ultrasound (24). The anchor-pulling condition *in vivo* on the other hand (*SI Appendix*, Fig. S6 *J*), is accomplished differently. Neurons are embedded in extracellular matrixes (earthy yellow color) such as laminin, poly-lysine, or poly-ornithine in the brain. When ultrasound is applied to the brain, extracellular matrixes anchor the ASIC1a (earthy arrow) while cytoskeletal changes pull it in a different direction (purple arrow), causing an activation of the cells manifesting as ERK phosphorylation.

Since DCX has been accepted as a neurogenesis markers in dentate gyrus (14, 25), increased DCX-positive cells in dentate gyrus with a consecutive 3-day of ultrasound treatment suggests a therapeutic implication. Of note, DCX plays multiple roles in brain development including hippocampal pyramidal neuronal lamination, cortical neuronal migration and axonal wiring (13, 26). The use of ultrasound in developing brains should be extremely cautious.

In conclusion, here we provide evidence that a clinically safely low-intensity transcranial ultrasound could modulate neuronal activity in mouse brain. The low-intensity ultrasound can directly activate neurons via tether-mode mechanotransduction and ASIC1a, which provides a molecular basis for future development of ultrasound neuromodulation.

## ACKNOWLEDGEMENT

This study was supported by Ministry of Science and Technology, Taiwan (MOST 107-2221-E-002-068-MY3, MOST108-2321-B-002-047, MOST 108-2321-B-002-061-MY2), National Health Research Institute, Taiwan (NHRI-EX109-10924EI), National Taiwan University (NTU-CC-107L891105); and a grant from MOST, Taiwan (MOST 108-2321-B-001-028-MY2) to CCC.

