## Supplementary information appendix for "ASIC1a is required for neuronal activation via low-intensity ultrasound stimulation in mouse brain"

Jormay Lim, Ya-Cherng Chu, Chen-Ming Hao and Wei-Hao Liao contribute equally.

Address correspondence and requests for reprints to

Jaw-Lin Wang, Ph.D., corresponding author

Professor, Department of Biomedical Engineering, College of Medicine and College of Engineering, National Taiwan University,

Address: 602 Jen-Su Hall, 1 Section 4, Roosevelt Road, Taipei 10617, Taiwan, ROC

Chih-Cheng Chen, Ph.D. corresponding author

Research fellow, Institute of Biomedical Sciences, Academia Sinica

Address: 128 Academia Road, Section 2, Taipei 115, Taiwan

**Supplementary materials**

**Methods and materials**

**Ultrasound devices and stimulation parameters**

Two different ultrasonic setups were used in our study. We used a commercial 1MHz transducer (C539-SM, Olympus, Tokyo, Japan) for mouse brain stimulation in the *in vivo* experiment (Fig. 1A). Simulation for calcium imaging was done with micropipette ultrasound attaching a 1MHz transducer (15 mm in diameter). Schematics of experimental setup for ultrasound calcium imaging is shown in Fig. 2A. All the transducers were controlled by a function generator (Tektronix AFG1022, Beaverton, OR, USA) through a power amplifier (E&I 210L, Rochester, NY, USA). The input voltage was 900 mVpp with a duty factor of 1% at 1 kHz pulse rate for the *in vivo* stimulation. We characterized cellular exposure to ultrasound using a hydrophone (HGL-1000, Onda, Sunnyvale, CA, US) immersed in water. The intensity used for in vivo animal experiments was 5 mW/cm$^2$ ($I_{SPTA}$), and 7.4 mW/cm$^2$ ($I_{SPPA,}$ at 700 mVpp) for micropipette in vitro experiment. These intensity values are within the range of not causing any side effects from ultrasound, like heat and cavitation.

Exploring upstream mechanoreceptors requires a calcium imaging assay that can respond to ultrasound stimulation repeatedly and reliably so that the effect of inhibitors can be demonstrated clearly. Micropipette ultrasound offers a wide range controllability. To select appropriate parameters for calcium experiments, we tested two extreme conditions: one with high input voltage and low duty factor (1500 mVpp, 0.05% duty factor) for a predominant ultrasound stimulus and the other with low input voltage (100 mVpp) and continuous waves for a predominant acoustic streaming stimulus. As a predominant ultrasound stimulus, micropipette ultrasound exhibit a point source characteristic (SI Appendix Fig. S4A). As a predominant streaming stimulus, micropipette ultrasound yields an inward flow pattern (SI Appendix Fig. S4B).

**Animals**

All animal procedures complied with the guidelines of the Institutional Animal

Care and Use Committee in Academia Sinica, Taipei, Taiwan. Either wildtype or ASIC1a$^{-/-}$ C57B6/J mice of 6-8 weeks were shaved under isoflurane anesthesia the day previous to ultrasound stimulation. The mice were randomly assigned to be either sham treated by placing ultrasound transducer on top of head or really exposed to ultrasound for 1 minute to evaluate neuronal activities in the mouse brain after the ultrasound stimulation under isoflurane anesthesia. Immediately after the treatment, mice were first anesthetized with urethane (1.5g/ kg; intraperitoneal) and perfused transcardially with 25 ml 0.02M Tris buffer saline (1x TBS, pH7.4, at 4°C) and then 25 ml cold fixative (4%[w/v] formaldehyde, 0.02M TBS (pH7.4, at 4°C).

**Brain histology and immunohistochemistry**

Mouse brain was dissected and post-fixed with 4% formaldehyde at 4°C for 16 hours; tissues were sectioned with Vibratome 1000 Plus (Rankin Biomedical, Holly, MI) at 100 µm thickness and incubated with antibody in free-floating method. For ABC-DAB-Nickel staining, tissue sections were first bleached in 1x TBS containing 0.03% $H_2O_2$ for 30 minutes, and then blocked in TBST (TBS+0.05% Triton X-100) containing 5% bovine serum albumin (BSA) (Sigma-Aldrich, St. Louis, MO, USA) and 5% normal goat serum (NGS from Jackson ImmunoResearch Laboratories, West Grove, PA, USA) at room temperature for 60 mins, and incubated with Rabbit polyclonal Phospho-p44/42 MAPK (ERK1/2) (Thr202/Tyr204) primary antibody [(1:500) #9101, Cell Signaling Technology, Danvers, MA, USA] diluted in blocking solution overnight at 4°C. Sections were then washed 3 times with TBST and incubated with secondary biotinylated goat-anti-rabbit antibodies (1:1000, Vector Laboratories, Burlingame, CA, USA) for 1 hour at room temperature. After 3 TBST washes, sections were incubated in the Avidin-Biotin pre-mix solution (1:200, Vector Laboratories, Burlingame, CA, USA). After 3 1xTBS washes, positive immunoreactivity signals were visualized using a Nickel-DAB method [DAB Peroxidase (HRP) Substrate Kit (with Nickel), 3,3’-diaminobenzidine SK-4100, Vector Laboratories, Burlingame, CA, USA or Sigma-Aldrich, St. Louis, MO, USA].

**Primary cell culture**

In order to ensure the detection of neuron specific p-ERK, we set up primary culture from neonatal mouse brain. Briefly, cortex isolated from neonatal mouse brain were mechanically minced by Castroviejo scissor and trypsinized by Trypsin (SI-T4174-100ml, Thermo Fisher Scientific, Waltham, MA, USA) diluted in Hanks Buffered Salt Solution (HBSS) (SI-H6648-500ml, Thermo Fisher Scientific, Waltham, MA, USA) with L-glutamine (2 mM/ml) (SI-G7513-100ml, Thermo Fisher Scientific, Waltham, MA, USA) for 15 minutes at $37^0$C with 3 subsequent HBSS washes before treated by deoxyribosenuclease I (SI-D4513-1vl, Thermo Fisher Scientific, Waltham, MA, USA). The treated tissues were then triturated with fire polished glass pipette and strained through 40[illegible]m strainer (431750, Corning Inc., Corning, NY, USA) and seeded on plasma treated and poly-D-Lysine (SI-P7405 Thermo Fisher Scientific, Waltham, MA, USA) coated glass cover slips at a density of $10^5$/ml in B27+ supplemented (Gibco A3582801, Thermo Fisher Scientific, Waltham, MA, USA) neurobasal media (Gibco A3582901, Thermo Fisher Scientific, Waltham, MA, USA) with 10% horse serum (Gibco 26050070, Thermo Fisher Scientific, Waltham, MA, USA) and penicillin/streptomycin (100 U/ml) (Life Technologies, Carlsbad, CA, USA). Culture was gradually replaced with serum free B27+ neurobasal media until day 7 for either immunofluorescence or for live cell calcium signal detection.

**Live cell calcium signal imaging**

In order to visualize calcium signal in the neurites and in the cell bodies of neuron, we treated the primary cultures with two different green fluorescent dyes, i.e. Invitrogen™ Oregon Green™ 488 BAPTA-1, AM cell permeant (O6807, Thermo Fisher Scientific, Waltham, MA, USA) and Invitrogen™ Fluo-4, AM, FluoroPure™ grade (F23917, Thermo Fisher Scientific Waltham, MA, USA) respectively. Living primary culture on cover slip was immersed in HHBS (20mM Hepes pH7.4, 1mM $CaCl_2$, 0.5mM $MgCl_2$, 0.4mM $MgSO_4$-$7H_2O$, 5mM KCl, 0.4mM $KH_2PO_4$, 4mM $NaHCO_3$, 138mM NaCl, 0.3mM $Na_2HPO_4$, 6mM D-Glucose) with 2-5 μM of fluorescent dye and incubate in incubator for 90 minutes. Subsequently, calcium staining solution was replaced with HHBS with 17% neurobasal media. Cover glass was mounted on an imaging chamber and placed under the fluorescent microscope and micropipette ultrasound was set up to the proximity of targeted cells.

Images were recorded using Olympus IX71 fluorescent microscope (Olympus Corporation, Shinjuku, Tokyo, Japan) with digital camera for microscope Camera attachment with 0.63x lens (DP80, Olympus Corporation, Shinjuku, Tokyo, Japan). Stacked images were analyzed in ImageJ. ROI of neurites or cell bodies were determined for stacks resliced to obtain data of fluorescence intensities plotted against time points.

**Molecular signaling protein inhibitors**

To determine whether PIEZO receptor or TRPC1 was responsible for the signal, we applied the GsMTx-4 (500nM) (1, 2) (ab141871, Abcam Inc., Cambridge, MA, USA) isolated from tarantula venom and Gadolinium (10 μM) (3) (G7532, Sigma-Aldrich, St. Louis, MO, USA) to the tissues or cells before ultrasound treatment. To investigate the potential role of ASIC channels in ultrasound signal transduction, we utilized the inhibitors such as Amiloride (100 μM) (4) (A7410-1G Sigma-Aldrich, St. Louis, MO, USA) and PcTx1 (0.1-50nM) (5) (Tocris #5042, Bio-Techne Corporation,

Minneapolis, MN, USA). To test whether endoplasmic reticular stored calcium was involved in the calcium signal detected, RyR inhibitor, JTV519 fumarate (10 μM) (6) (Tocris #4564, Bio-Techne Corporation, Minneapolis, MN, USA) and Thapsigargin (T9033, Sigma-Aldrich, St. Louis, MO, USA) was tested.

**Immunofluorescence Staining of DCX**

After the VLIUS stimulation, the mice were sacrificed and perfused with 10% formaldehyde/PBS. The brain was then harvested and fixed with 10% formaldehyde/PBS at room temperature. Samples were embedded in paraffin and serial 7μm transverse sections were mounted on slides. The samples were deparaffinized, rehydrated, antigen retrieved (100℃, 20 min) and washed in PBST. Slices were blocked with 10% newborn calf serum (NCS) and 1% BSA in PBST for 1 hours, incubated with primary antibody overnight at 4℃. After washing with PBST, the samples were incubated with the secondary antibody for 1 hours at room temperature, washed with PBST and mounted with EverBrite™ Hardset Mounting Medium containing DAPI to label the nuclei (Biotium). Slides were viewed, and images were captured with LSM780 confocal microscope (Zeiss, Jena, Germany). The primary antibodies used for immunostaining and their dilutions were as follows: rabbit anti-DCX (1:200, Cell signaling), mouse anti-MAP2 (1:200, Thermo). The secondary antibodies used were Alexa Fluor 488-conjugated goat anti-rabbit IgG (1:100, Thermo) and Alexa Fluor 555-conjugated goat anti-mouse IgG (1:100, Thermo).

**Data and Statistical Analyses**

Cells were counted using ImageJ. Cells were identified using a global threshold with watershed segmentation. The number of pixel groups was evaluated as the number of cells. Cells were also manually counted from bright-field images. Measurements were compared between control and ultrasound groups using student t-test. A *p* value $\leq 0.05$ was considered to indicate statistical significance. All statistical analyses of animal studies were performed using GraphPad Prism 8.

Supplementary Table 1: In vivo animal and human transcranial ultrasound experiments.

| Reference | Disease & Model | Ultrasound parameters | Finding |
|---|---|---|---|
| Liu 2019 (7) | Acute ischemic stroke, In vivo rat model | 0.5 MHz, Isppa=2.6 W/cm2, (Ispta=173 mW/cm2), Dose: 10 min/day | Ultrasound increase the Apparent Diffusion Coefficient (ADC) of MRI, the earlier intervention (0.5 h) is better than late (up to 9 h).<br>Rationale: US increase fluid flow within the brain |
| Eguchi 2018 (8) | Vascular & Alzheimer's Dementia, In vivo mice model | 1.875 MHz, DF=10%, Ispta=90 mW/cm2, Dose: 20 min *3 days | LIPUS improved cognitive dysfunctions due to eNOS expression.<br>Rationale: US increase the vascular endothelial cell eNOS. |
| Sato 2016 (9) | Inferior alveolar nerve (IAN) injury, In vivo rat model | 1 MHz, Isata=30 mW/cm2, Dose: 20 min/day for 28 d | Increased mechanical sensitivity, and TG cell number.<br>Rationale: US increase trigeminal ganglion (TG) cell number. |
| Baek 2018 (10) | Cerebellar ischemic stroke, MCAO model, in vivo mice model | 0.35 MHz, Isppa=2.54 W/cm2, (Ispta=127 mW/cm2), Dose: 20 min/day for 2 d | Increased ipsilateral water content due to tissue swelling, showing attenuation of brain edema. Prominently, the reduction of neuro-immune reactivity at the infarct core and peri-infarct region. |
| Lin 2015 (11) | Al-induced Alzheimer dementia, In vivo Rat model | 1 MHz, DF=5%, Ispta=528 mw/cm2, Dose: 5 min*3/day | Increase BDNF, GDNF, VEGF, Decrease Al particle, acetylcholinesterase, beta-amyloid, karyopyknosis, Increased behavioral test.<br>Rationale: US increase BDNF |
| Hung 2017 (12) | Bilateral common carotid artery occlusion (BCCAO) induced vascular dementia | Same as above, | Increased BDNF, myelin using micro-PET images, and histology Recovered hippocampus neuron, increased behavioral test<br>Rationale: US increase BDNF |
| Su 2017 | Cortical impact | Same as above, | Reduced brain edema, blood |

| | | | |
|---|---|---|---|
| (13), Chen 2018 (14), Su 2017 (15) | injury induced brain trauma | | brain barrier permeability, and neuronal degeneration at day 1, improved functional recovery and reduced contusion volume at day 28. At day 4 reduced MMP9, increased BDNF, enhance p-TrkB, Akt, cAMP response<br>Rationale: US increase BDNF |
| Chen 2018 (16) | Cerebral ischemia/reperfusion injury using middle cerebral artery occlusion (MCAO) model | Same as above | Apoptosis reduction and BDNF induction in MCAO model<br>Rationale: US increase BDNF |
| Chen 2019 (17) | Lipopolysaccharide (LPS) induced Alzheimer's | Same as above | Increased behavioral test, Decreased beta-amyloid, APP, Caspase-3, GFAP, TNF-alpha, IL-1beta, IL-6 at hippocampus, and cortex, NF-kapaB, TLR4 signal pathway, Increased BDNF and CREB<br>Rationale: US increase BDNF |
| Lin 2018 (18) | Brain glioblastoma | 1 MHz, A392S transducer Energy= 2.86 W, DF=5%, Stimulation: 60 sec | US as a BBB disruption, increased Dox concentration |
| Tyler 2008 (19) | mice brain slice (400 micron) | 0.44 MHz, Isppa=2.9 W/cm2, Ispta=23 mW/cm2, DF=1%, P=0.8MPa, | US regulate voltage-gated sodium and calcium channels. Trigger SNARE-mediated exocytosis and synaptic transmission. |
| Nicodemus 2019 (20) | Clinical trial of focused US for AD | Ispta=520 mW/cm2, DF=0.1%, P= 5 MPa, Stimulation=1 hour | Cognition: 1/3 improved, 1/3 decreased, 1/3 remain same<br>Motor: 5% improved, 15% decreased,<br>MRI showed immediate increased blood flow perfusion. |
| Legon 2014 (21), Panczykowski 2014 (22) | Human cortical | 0.5 MHz, Isata=4.295 W/cm2, DF=36%, P=0.8MPa, Stimulation=0.5 sec | Ultrasound can focally modulate cortical function, indicated by changed alpha, beta, gamma wave function |

Supplementary Table 2. Immunohistochemical staining of p-ERK expression in Control and Ultrasound treated wild type mouse brain

| Brain region (Bregma:-1.5,-2) | WT Control | Ultrasound |
|---|---|---|
| **Cortex (Posterior parietal association areas)** | **-** | **+** |
| **Cortex (Motor area)** | **-** | **-** |
| - Secondary motor area | - | - |
| - Primary motor area | - | - |
| **Cortex (Somatosensory area)** | **+** | **+** |
| - Primary somatosensory area, trunk | - | + |
| - Primary somatosensory area, barrel | - | + |
| **Cortex (Auditory area)** | **-** | **+** |
| - Dorsal auditory area | - | + |
| - Primary auditory area | - | + |
| - Ventral auditory area | - | + |
| - Temporal association areas | - | + |
| **Cortex (Entorhinal area)** | **+** | **++** |
| **Endopiriform nucleus** | **+** | **-** |
| **Caudoputamen** | **-** | **+** |
| **Hippocampus (CA1 subfield)** | **-** | **-** |
| **Hippocampus (CA2 subfield)** | **-** | **-** |
| **Hippocampus (CA3 subfield)** | **-** | **-** |
| **Hippocampus (dentate gyrus)** | **-** | **-** |
| **Central Amygdalar nucleus** | **-** | **+++** |
| **Basolateral Amygdalar nucleus** | **+** | **+** |
| **Cortical Amygdalar nucleus** | **-** | **+** |
| **Medial Amygdalar nucleus** | **-** | **+** |
| **Piriform cortex** | **+** | **++** |

| Brain region (Bregma:-2.7,-3) | WT Control | Ultrasound |
|---|---|---|
| **Retrosplenial area** | - | + |
| **Cortex (Visual area)** | - | + |
| - Anteromedial visual area | - | + |
| - Primary visual area | - | + |
| - Amterolateral visual area | - | + |
| **Cortex (Posterior parietal association areas)** | + | + |
| **Cortex (Auditory area)** | - | ++ |
| - **Dorsal auditory area** | - | + |
| - **Primary auditory area** | - | ++ |
| - **Temporal association areas** | - | + |
| **Cortex (Ectorhinal area)** | - | ++ |
| **Cortex (Piriform area)** | - | - |
| **Cortex (Entorhinal area)** | - | + |
| **Cortex (Perirhinal area)** | - | + |
| **Postpiriform transition area** | ++ | + |
| **Cortical Amygdalar area** | - | - |
| **Hippocampus (CA1 subfield)** | + | ++ |
| **Hippocampus (CA3 subfield)** | - | - |
| **Hippocampus (dentate gyrus)** | + | ++ |
| **Superior colliculus** | - | + |
| **Subiculum** | - | + |

The sections were incubated with anti-p-ERK antibody and stained with avidin–biotin-peroxidase for visualization. p-ERK expression was quantified by the number of positive staining cells was calculated using optical microscopy in 100 μm coronal brain sections from control and ultrasound treated wild type mice brain. ‘‘-’’ absent; ‘‘+’’: low expression; ‘‘++’’ : moderate expression; ‘‘+++’’: high expression

**SI Appendix Figure captions:**

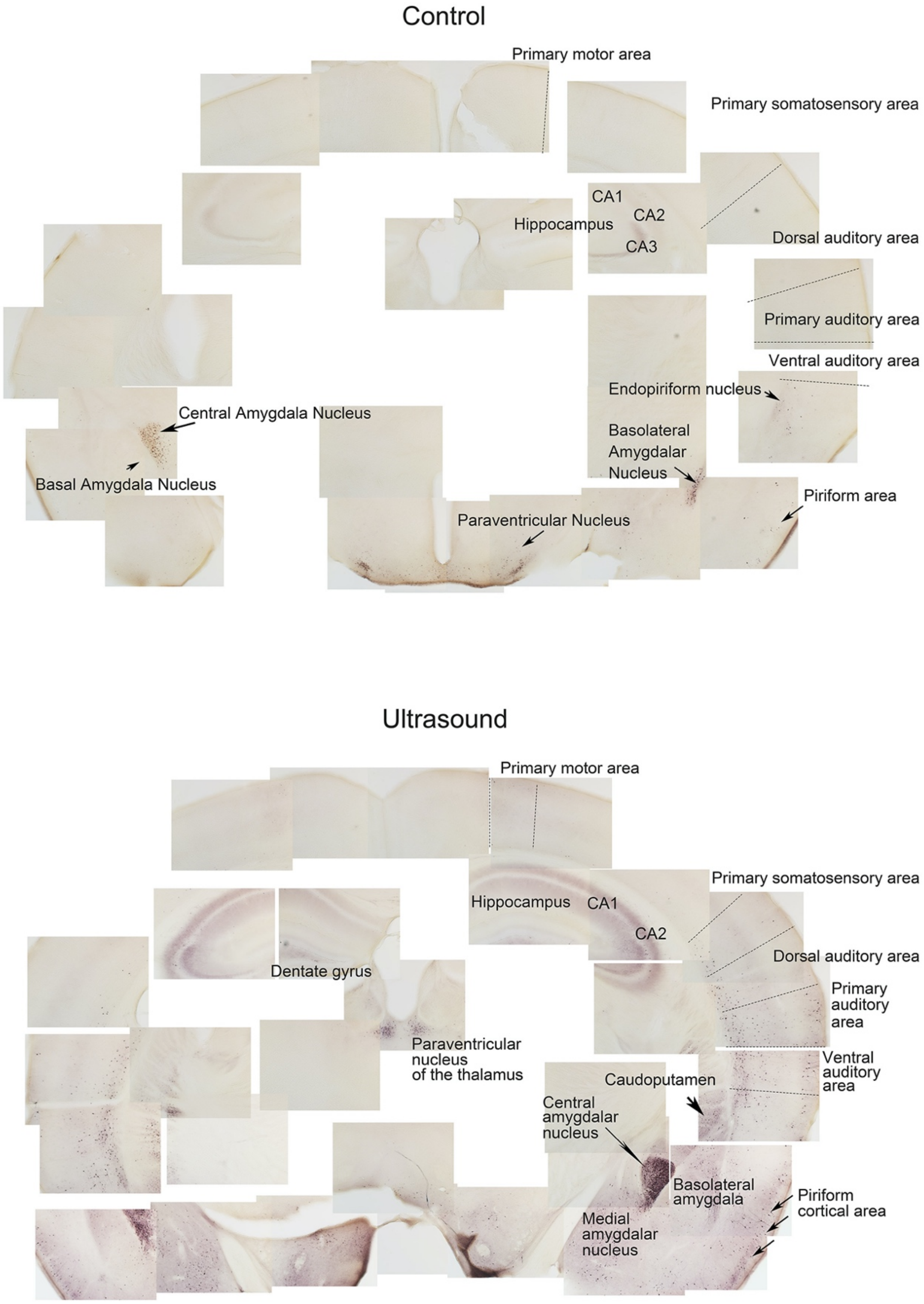


***SI Appendix,* Fig. 1**: Stitched images showing p-ERK expression of brain slice containing hippocampus, cortex and amygdala of sham control v.s. ultrasound stimulated mice.

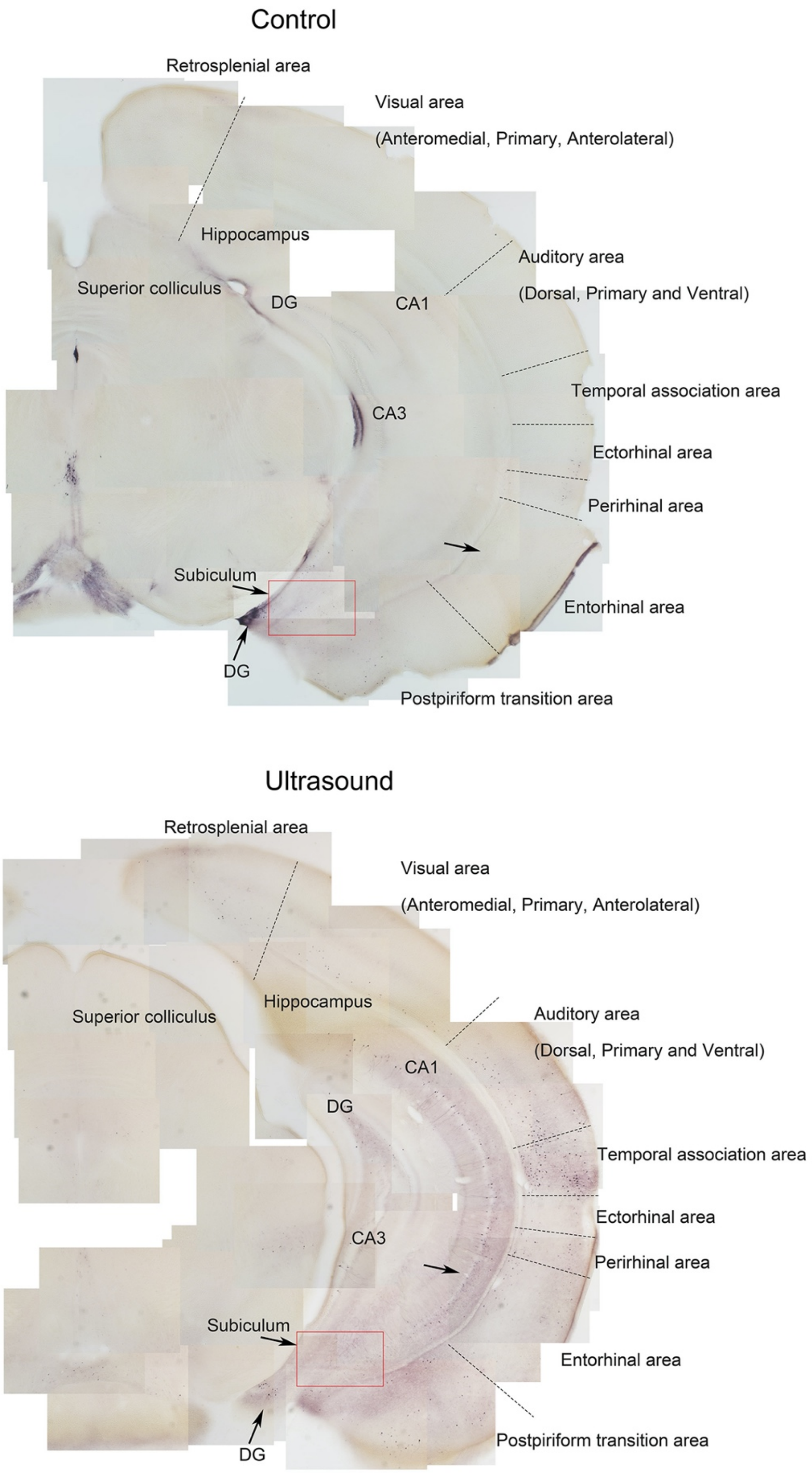


***SI Appendix,* Fig. 2**: Stitched images showing p-ERK expression of hippocampus and piriform cortex of sham control v.s. ultrasound stimulated mice.

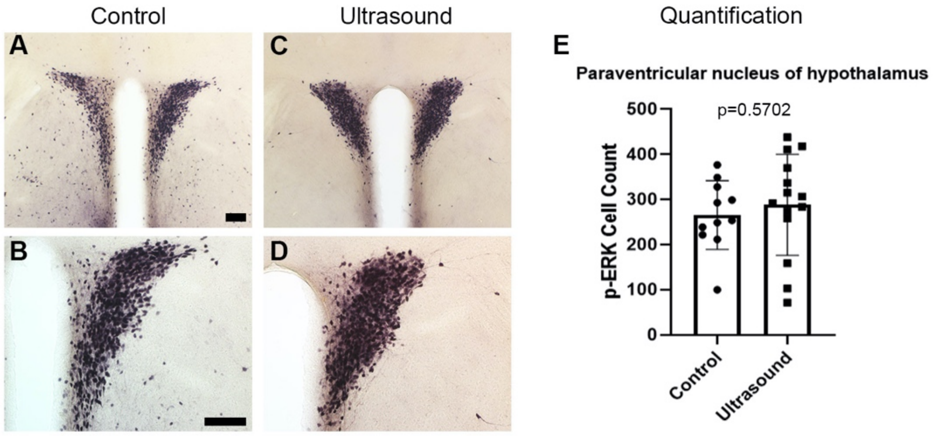


***SI Appendix,* Fig. 3**: The similar p-ERK expression of paraventricular nucleus of hypothalamus (PVH) of control untreated vs ultrasound stimulated mice.

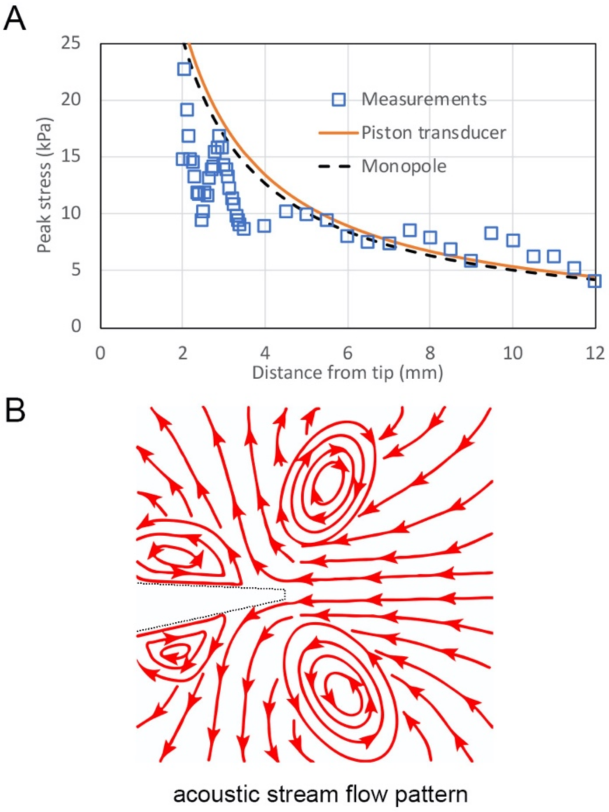


***SI Appendix,* Fig. 4**: (*A*) The measured and monopole modeled peak pressure stress vs distance from tip of micropipette. (*B*) The schematic acoustic stream flow pattern depicted from Supplementary Movie 1.

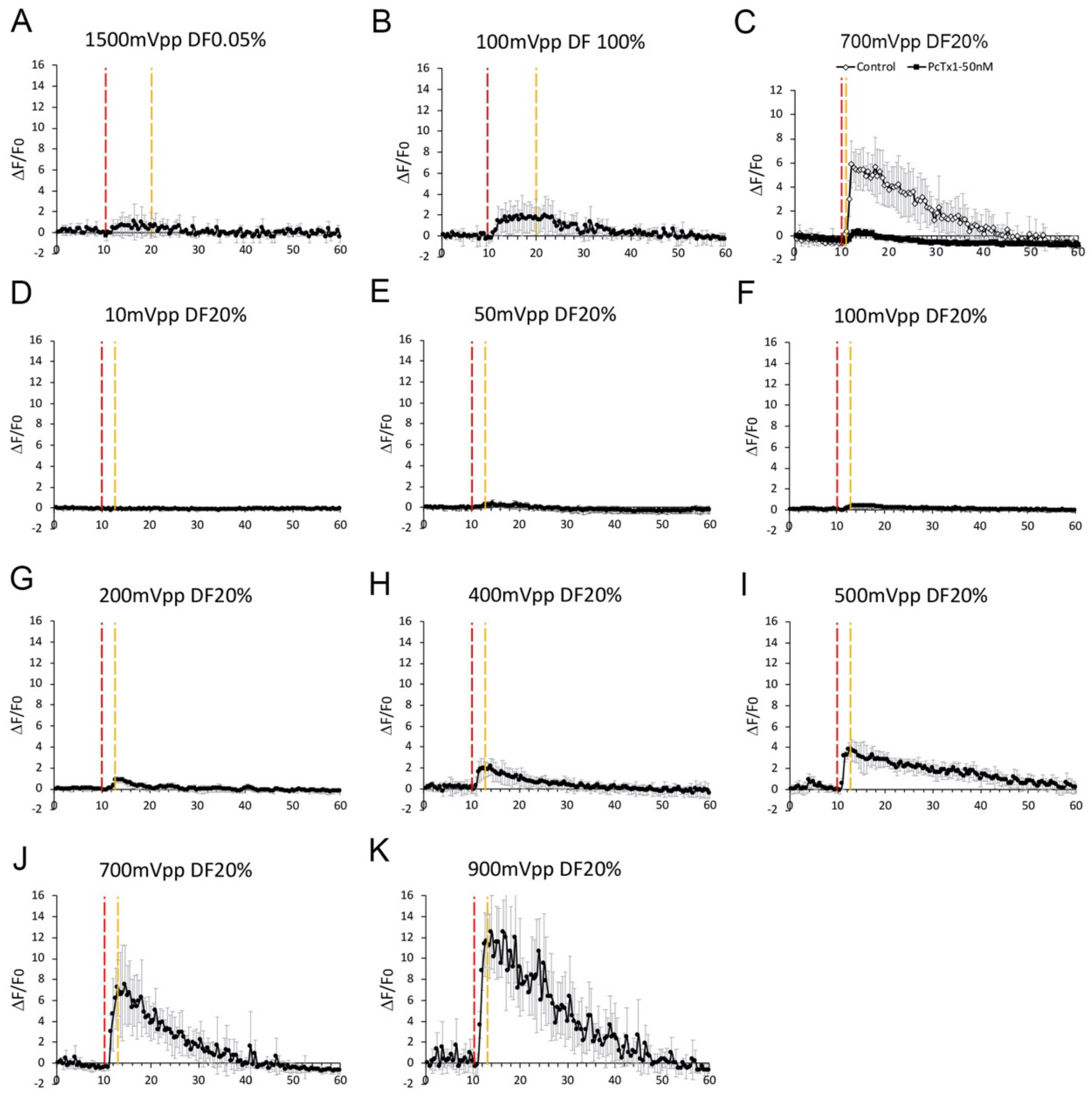


***SI Appendix,* Fig. 5**: (*A*) The ultrasound induced calcium response of 1500mVpp at 0.05% duty factor for 10 seconds was minimal. The physical effect of pressure stress is dominant in this stimulation condition. (*B*) The ultrasound induced calcium response of 100mVpp at 100% duty factor for 10 seconds was also minimal. The physical effect of acoustic streaming is dominant in this stimulation condition. (*C*) The elevated ultrasound induced calcium response of 700mVpp at 20% duty factor for as short as 1 second. The response was diminished by applying PcTx1 (50nM), an ASIC1a inhibitor. (*D*-*K*) The dose response of ultrasound induced calcium response from 10 mVpp to 900 mVpp at 20% duty factor. The ultrasound stimulation time were 3 seconds.

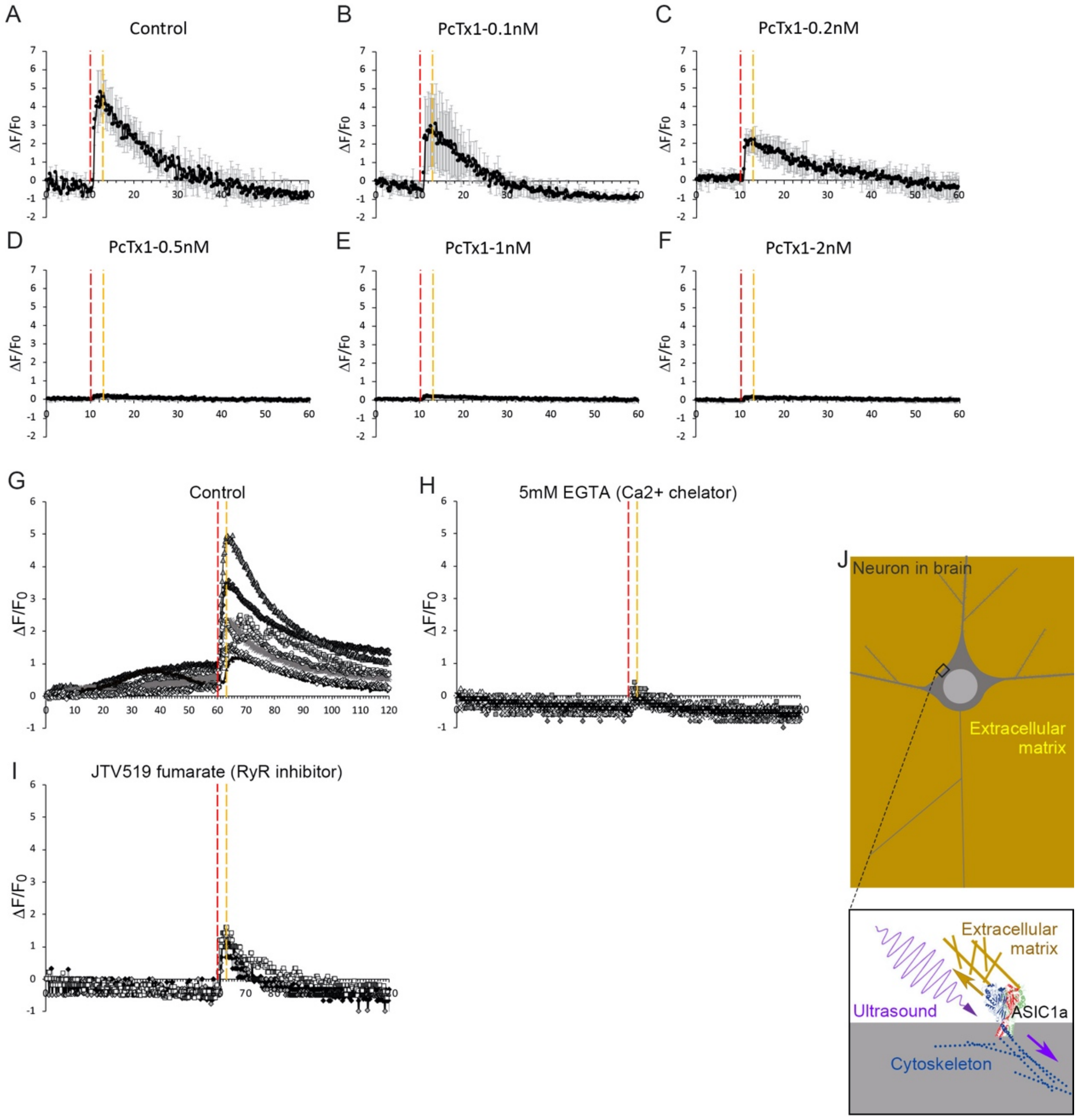


***SI Appendix,* Fig. 6**: (*A*-*F*) The dose-dependent calcium response of PcTx1, an ASIC1a inhabitation, under micropipette stimulation (700mVpp, 20%DF, check). The ultrasound stimulation time was 3 seconds. (*G*) This figure is not used in main manuscript. (*H*) The EGTA (a calcium chelator) inhibit the calcium response of ultrasound stimulation (5 mM). (*I*) The JVT519 fumerate, a RyR inhibitor, also moderately inhibit calcium response of ultrasound stimulation (dose). (*J*) A schematic tether mode mechanotransduction model for in vivo circumstance.
